# *In vivo* validation of predicted fitness effects at single-base resolution in a *Brachypodium distachyon* mutant population

**DOI:** 10.64898/2026.03.31.715642

**Authors:** Camous Moslemi, Marine Folgoas, Xiaqing Yu, Julie D. Jensen, Stephan Hentrup, Tianyi Li, Hai Wang, Birte Boelt, Torben Asp, Richard Sibout, Guillaume P. Ramstein

**Affiliations:** Center for Quantitative Genetics and Genomics, Aarhus University, 8000 Aarhus, Denmark; Department of Clinical Medicine, Aarhus University, 8200 Aarhus, Denmark; Earlham Institute, NR4 7UZ Norwich, United Kingdom; College of Horticulture, Nanjing Agricultural University, 210095 Nanjing, China; Department of Agroecology, Aarhus University, 4200 Slagelse, Denmark; Department of Plant Genetics and Breeding, China Agricultural University, 100193 Beijing, China; INRAE, UR1268 BIA, F-44316, Nantes, France

**Keywords:** Variant effect prediction, Biological language models, Sequence-to-function models, Untargeted mutagenesis, Brachypodium distachyon

## Abstract

Computational tools, including biological language models (LMs), show substantial promise in predicting the impact of genetic variants on plant fitness. However, validating variant effect predictions (VEP) requires experimental populations where genetic variation consists of discrete point mutations rather than segregating recombination blocks. In this study, we generated a novel population of *Brachypodium distachyon* mutant lines to evaluate the accuracy of VEP at single-base resolution. These lines were advanced through single-seed descent for five generations (M_1_ to M_5_), with whole-genome sequencing performed at M_2_ and M_5_ and phenotypic measurements recorded at M_3_ and M_4_. Using state-of-the-art VEP models, we predicted the functional impact of missense protein-coding variants and gene-proximal non-coding variants. We validated these predictions by estimating the effect of mutations on whole-plant measurements (burden tests) and their probability of fixation from M_2_ to M_5_ (purging tests). Among missense variants, the protein LM ESM showed superior predictive accuracy compared to the bioinformatic standard SIFT and the genomic LM PlantCAD. Notably, the relationship between VEP scores and allele fixation suggested a log-linear relationship between VEP scores and variant fitness. Among gene-proximal variants, PlantCAD appeared more accurate than supervised models of regulatory activity, such as chromatin accessibility (a2z) and RNA abundance (PhytoExpr). Collectively, our findings highlight the utility of state-of-the-art VEP tools as predictors of fitness and demonstrate the potential of mutant populations to evaluate computational tools for precision breeding applications.

## INTRODUCTION

In plant breeding, recent computational advances in variant effect prediction (VEP) provide a unique opportunity for the efficient identification of impactful mutations as targets for selection or genome editing (1). Of particular interest is the prediction of variant effects on plant fitness, which allows breeders to target alleles with potential benefits for traits such as crop yield and resilience (2). Among the most promising developments in this area are sequence-based deep learning techniques, specifically biological language models (LMs) and sequence-to-function models.

Biological LMs evaluate the fitness effects of point mutations by estimating their likelihood within a specific protein or DNA sequence context (3, 4). ESM (Evolutionary Scale Modeling) has emerged as one of the most effective protein LMs for scoring the fitness consequences of protein-coding variants (3, 5). Despite being trained on a massive cross-species corpus (UniRef90), ESM has demonstrated high VEP accuracy in studies focused on human and *in vitro* assays (6–8). Notably, ESM has consistently outperformed SIFT (Sorting Intolerant from Tolerant), a traditional bioinformatic tool that relies on multiple sequence alignments (MSAs) to assess evolutionary conservation of amino acid residues (9).

Genomic LMs extend fitness effects predictions beyond protein-coding regions to include non-coding variants (4). Unlike protein LMs, which analyze whole protein sequences, genomic LMs typically predict variant effects within a fixed-length DNA sequence window. PlantCaduceus (PlantCAD) is a prominent example; trained on the genomes of 16 angiosperm species, it has proven effective at VEP in both coding and non-coding regions, as validated by allele frequency data across diverse accessions (10). Genomic LMs hold significant potential for VEP, because they are inherently versatile and, critically, do not require gene annotations to score variants.

A distinct category of sequence-based deep learning involves sequence-to-function models, which use DNA sequences to predict molecular phenotypes such as chromatin accessibility (11) or RNA abundance (12). The regulatory effects of variants can be predicted by chromatin-state models like a2z, which was trained on ATAC-seq data from 12 angiosperms to predict the impact of DNA variants on chromatin accessibility in leaf tissues (11). While chromatin accessibility is a secondary proxy for fitness, it has been shown to affect gene regulation (13) and frequently co-occur with causal variants (14–16).

The regulatory impact of variants can also be assessed via sequence-to-expression models. Although benchmark studies in humans suggest that chromatin-state models may currently be more effective for VEP than sequence-to-expression models (17), it remains unclear whether the same is true in plants. In plants, the recent PhytoExpr model (12) has achieved high accuracy in predicting mRNA abundance from *cis*-regulatory sequences across 17 species. Similar to previous architectures (18, 19), PhytoExpr focuses on gene-flanking sequences, specifically analyzing sequences spanning 4 kb upstream of the transcription start site (TSS) to 1 kb downstream, and 1 kb upstream of the transcription termination site (TTS) to 4 kb downstream. As demonstrated by studies of chromatin accessibility across angiosperms, these ranges are expected to capture the majority of *cis*-regulatory features (15).

Despite the rapid development of VEP techniques, their empirical validation for predicting fitness effects in plants still lags behind (1). Most current VEP benchmarks in this field rely on natural accessions (10, 20, 21), where associations between variants and phenotypes are confounded by extensive linkage disequilibrium (LD) and historical selection (22, 23). Furthermore, few studies have systematically compared the performance of LMs and sequence-to-function models within the same experimental context. To address these limitations, we developed the SIEVE (Selection of mutations by *in silico* and experimental variant effects) population through untargeted mutagenesis in *Brachypodium distachyon*.

Untargeted mutagenesis provides a unique opportunity to observe the phenotypic impacts of point mutations with minimal confounding from LD, provided that the mutagenized lines remain independent (i.e., they have shared no common haplotypes since the mutagenesis event). While mutant populations have previously been generated in *B. distachyon* and related taxa, these have primarily been used for candidate-variant analysis by targeted genotyping (e.g., TILLING) (24–26). To our knowledge, none have been employed for the systematic validation of VEP models. In this study, we used sodium azide (NaN_3_) to generate point mutations – predominantly G-to-A and C-to-T transitions – and evaluated their effects across independent lines derived from a single seed source. This experimental design enabled us to assess the impact of variants prioritized by VEP techniques at single-base resolution rather than LD-block resolution in natural populations.

We conducted our study in the model grass *B. distachyon* because of decisive advantages: its compact genome (272 Mb), small stature, naturally selfing habit, and rapid generation time (27), as well as its close phylogenetic proximity to major cereal crops like wheat, barley, and rice (28). We aimed to evaluate the performance of leading VEP techniques for predicting effects of induced variants in protein-coding regions (SIFT, ESM, PlantCAD) and non-coding regions flanking the genes’ TSS and TTS (a2z, PhytoExpr, PlantCAD). We incorporated predicted variant effects into statistical models to assess their association with whole-plant phenotypes (e.g., total seed weights) and allele fitness (measured as the probability of fixation across generations). As demonstrated here, the SIEVE population constitutes a valuable benchmark for current and future VEP techniques, especially for validating and calibrating predicted fitness effects at single-base resolution.

## Material and methods

### Plant material

Seeds from *Brachypodium distachyon* Bd21-3 were produced by two generations of selfing from a unique seed (provided by R. Sibout, INRAE). At the M_1_ generation, four batches of seeds were sown. In each batch, 2080 seeds were exposed to 7 mM of NaN_3_, while 104 control seeds were exposed to the same solution with no mutagen. Treated seeds were then sown in trays in the greenhouse (20h/4h light/dark, 21°C/18°C target temperature, at Aarhus University Flakkebjerg). Successfully grown seedlings were then transferred to pots: 1808 mutants and 96 controls at generation M_1_. Pots from each batch were arranged in two ‘blocks’, each of which corresponded to one half of a table (eight blocks and 4 tables used to grow all plants at M_1_). The physical location of blocks changed from one generation to the next, but plants were assigned to the same blocks across generations.

At each generation from M_1_ to M_5_, plants were allowed to self-fertilize naturally to produce the next generation. From generation M_2_ to M_5_, seeds from each plant line were pre-stored for one week at 37°C and 20% relative humidity, to improve and synchronize germination ^(^Barrero et al., 2012). At generations M_2_ and M_5_, leaf samples were collected from each plant line at the juvenile stage (about 5 weeks after sowing). At generations M_3_ and M_4_, each plant was phenotyped for the following traits: plant height (distance between the base of the plant and the tallest spike), germination rate (number of seeds that germinated out of 5), seed weight (total weight of harvested seeds), and heading date (number of days until about half of tillers reach stage 50 “First spikelet of inflorescence just visible” on the Zadoks scale) (29).

### Genotyping

At the M_2_ and M_5_ generations, juvenile leaf tissue was collected (about 150 mg of leaf tissue, 5 weeks after sowing), for subsequent DNA extraction and whole-genome sequencing by DNBSEQ (paired-end sequencing, 150-bp reads) at BGI, Hong-Kong. Sequencing reads (fastq files) were aligned to the Brachypodium distachyon Bd21-3 v1.2 assembly from Phytozome (BdistachyonBd21_3_537_v1.0.fa), by BWA-MEM (30). Then, SNP variants were called by GATK (31). Finally, variant sites were filtered based on GATK’s standard filters: QD < 2, QUAL < 30, SOR > 3, FS > 60, MQ < 40, MQRankSum < -12.5, ReadPosRankSum < -8. The resulting VCF file contained genotype calls for 1998 individuals and 1,046,483 variants.

To estimate relatedness among individuals due to naturally occurring variants and potential accidental outcrossing, the genomic relationship **G** was calculated by the R package SNPRelate (32), following (33).

Since plant genomes descended from the same seed of the reference genotype Bd21-3, singleton variants were likely induced by mutagenesis. Moreover, variants induced by NaN_3_ are generally G:C-to-A:T transitions (hereafter, ‘GC-transitions’) (24). Therefore, variants were called induced if they met the following criteria: (*i*) only one occurrence of the alternate allele at heterozygous or homozygous state among all genotyped samples at M_2_; and (*ii*) the reference allele being G or C and the alternate allele being the corresponding transition (A or T). Singleton variants were further selected based on the genotype quality (GQ) score at their unique variant genotype: GQ ≥ 20 at the M_2_ and M_5_ generations (filtering out false variant calls due to sequencing or genotyping errors).

To ensure that the study sample was not affected by sample mix-ups, accidental outcrossing, or genotyping errors, mutant lines were discarded if they failed to meet all of the following criteria: genomic relationship below 0.1 with any other lines at M_2_, from M_2_ to M_5_, and at M_5_ (filtering out potential outcrossing lines); > 90% of singletons observed at M_5_ also observed at M_2_ (filtering out lines with residual genotyping errors at M_2_ and/or M_5_). After discarding lines based on these criteria, the resulting genomic data contained genotype calls for 920 lines (889 mutant lines and 31 control) and 786,305 selected singletons.

### Variant Effect Prediction

Variant effects were predicted for two categories of variants: (1) ‘missense variants’, causing amino-acid changes within protein-coding sequences; and (2) ‘gene-proximal variants’, comprising all variants included in PhytoExpr sequence-to-expression model ([-4 kb, +1 kb] from TSS, [- 1 kb, + 4kb] from TTS, excluding variants in protein-coding regions). For each annotated gene in the Bd21-3 genome, VEP scores were computed for the primary transcripts as defined by the ‘BdistachyonBd21_3_537_v1.2.transcript_primaryTranscriptOnly.fa.gz’ file in Phytozome (https://phytozome-next.jgi.doe.gov/info/BdistachyonBd21_3_v1_2).

#### SIFT scores

We used SIFT4G to generate SIFT annotations based on UniRef90, our VCF file, and the reference genome of *B. distachyon* (Organism: *Brachypodium distachyon* Bd21-3, NCBI-Taxonomy-ID: 15368 Assembly-version: v1.0, Annotation-version: v1.2) (https://github.com/pauline-ng/SIFT4G_Create_Genomic_DB). SIFT scores were then extracted from the SIFT4G database for each missense variant. These scores were values between 0 and 1 quantifying the probability of amino acid substitutions, such that low a SIFT score (e.g., below 0.05) suggests that the alternate allele is deleterious (34).

#### ESM scores

The SIFT4G database was used to identify missense variants. We then generated alternate protein sequences and used the five ESM-1v models to generate an average log-likelihood ratio score for each missense variant within the first 1022 residues (upper limit of ESM sequence size) of transcripts (https://github.com/facebookresearch/esm, esm1v_t33_650M_UR90S_[1-5].pt): 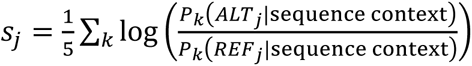, where *j* is a variant position, *P*_*k*_ is the predicted likelihood of the ALT or REF protein residue at position *j* based on the *k*th ESM-1v model (*k* = 1, …, 5), and ‘sequence context’ is the sequence of the focal protein, excluding position *j*.

#### PlantCaduceus scores

LLR scores of missense and gene-proximal variants were generated based on the PlantCaduceus l32 model (https://huggingface.co/kuleshov-group/PlantCaduceus_l32) and the zero shot scoring routine (https://github.com/plantcad/plantcad, zero_shot_score.py): 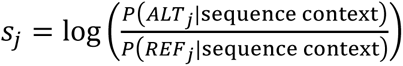, where *j* is a variant position, *P* is the likelihood of the ALT or REF nucleotide at position *j* predicted by the PlantCaduceus model, and ‘sequence context’ is the sequence of the 512-bp window centered at position *j*.

#### a2z scores

*In silico* mutagenesis (ISM) scores for chromatin accessibility were generated for each gene-proximal variant based on the full a2z model (https://zenodo.org/records/5724562, model-accessibility-full.h5): 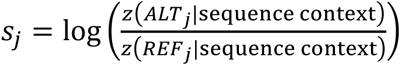, where *j* is a variant position, *z* is the predicted chromatin accessibility of the ALT or REF nucleotide at position *j*, and ‘sequence context’ is the sequence of the 600-bp window centered at position *j*.

#### PhytoExpr scores

ISM scores for mRNA abundance were generated for each gene-proximal variant based on the ensemble of five PhytoExpr models, each trained on a different cross-validation fold (12): 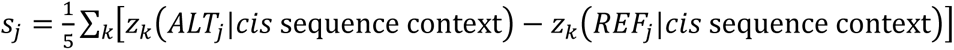, where *j* is a variant position, *z*_*k*_ is log(1+TPM) for the ALT or REF nucleotide at position *j* based on the *k*th PhytoExpr model (*k* = 1, …, 5), and ‘*cis* sequence context’ is the pair of upstream ([-4 kb, +1 kb] from TSS) and downstream ([- 1 kb, + 4kb] from TTS) sequences at the focal gene.

### Gene depletion analyses

Gene depletion analyses tested whether genes belonging to certain classes were less prone to carry induced (potentially deleterious) mutations, due to the genes’ contribution to plant fitness. Gene classes were pathway or gene ontology (GO) annotations.

Pathway annotations were obtained from PMN (Plant Metabolic Network) (35). The ‘brachypodiumcyc_pathways.20230103’ file was retrieved from the BrachyCyc database and used to obtain a set of genes within each annotated metabolic pathway.

GO annotations were obtained from the ‘BdistachyonBd21_3_537_v1.2.annotation_info.txt’ in Phytozome (https://phytozome-next.jgi.doe.gov/info/BdistachyonBd21_3_v1_2). The plant GO slim subset (https://www.geneontology.org/ontology/subsets/goslim_plant.obo) was then used to group similar GO annotations into 94 broad GO classes, excluding global classes GO:0003674 (molecular function), GO:0008150 (cellular component), GO:0009987 (biological process), and GO:0005575 (cellular process).

Only the gene classes containing more than 10 genes were included in these analyses.

#### Density of induced missense variants at M_2_ generation

The mutation density of each gene was calculated by using the SIFT database to identify all possible missense GC-transitions within a gene. The mutation density of any given protein was then calculated by dividing the number of observed missense GC-transitions (singletons) at the M_2_ generation by the total number of potential missense GC-transitions. For each gene class, we then fitted a logistic regression model (by the glm function in R), where the response was a binary indicator of whether each gene belonged to the gene class, and the predictors were the density of homozygous mutations and the density of heterozygous mutations in each gene.

#### Purging of missense variants across generations

To test whether some classes of genes were especially prone to purging induced mutations, we fitted a logistic regression model (by the glm function in R), where the response variable was the fate of each heterozygous variant from the M_2_ generation (‘purged’ if homozygous for the reference allele at M_5_, ‘fixed’ if homozygous for the alternate allele at M_5_, missing otherwise), and the predictors were a binary indicator of whether the variant belonged to the gene class and the number of singletons per chromosome arm as potential confounder.

To avoid heterogeneity among different categories of variants, only missense variants were included in this analysis. Moreover, only the gene classes represented by at least 20 variants were included in this analysis.

### Functional enrichment by VEP scores

#### Effect of prioritized variants on plant phenotypes (burden tests)

To test effects of induced mutations on plant phenotypes we used linear mixed models fitted by the R package MM4LMM (36) to correlate plant phenotypes with the number of induced singletons carried by each plant. To investigate whether VEP scores detected impactful variants, the effects of prioritized singletons were estimated in the following model:

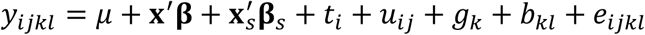

where *μ* is the intercept; **x** is the vector of mutational burdens (counts of alternate alleles at homozygous state, *x_hom_*, and heterozygous state, *x_het_*), **β** consists of their fixed effects, i.e., the average effect of alternate alleles at homozygous and heterozygous states; **x***_s_* is the vector of mutational burdens at prioritized variants (counts of alternate alleles, at homozygous and heterozygous states), **β***_s_* consists of their fixed effects; *t_i_* is the fixed effect of treatment *i* (control or mutant); *u_ij_* is the random genomic value of line *j* from treatment *i*, as captured by segregating variants, such that 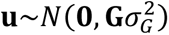, with **G** the genomic relationship matrix calculated from R package SNPRelate (32); *g_k_* is the fixed effect of generation *k* (M_3_ or M_4_); *b_kl_* is the effect of block *l* at generation *k*; *e_ijkl_* is the model residual. The effects of induced mutations were estimated based on genotypes at the M_2_ generation: homozygous at M_2_ (and therefore, likely homozygous at M_3_ and M_4_), or heterozygous at M_2_ and segregating at M_3_ and M_4_.

For a given VEP score and threshold (minimum or maximum value), only variants whose VEP score passed the threshold were included in **x***_s_* (i.e., they were prioritized only if their VEP score was above the minimum value or below the maximum value). Functional enrichment by VEP scores was then assessed by testing the significance of **β***_s_* estimates. Significance tests consisted of parametric tests (Wald tests) as well as permutation tests where the **β***_s_* coefficient estimates were compared to the null empirical distribution of estimates from 100 sets of permuted VEP scores.

#### Allele purging across generations (purging tests)

To investigate whether variants’ fixation probability depended on their VEP score, we fitted a logistic generalized additive model (GAM) on the fate of each heterozygous variant from the M_2_ generation (‘purged’ if homozygous for the reference allele at M_5_, ‘fixed’ if homozygous for the alternate allele at M_5_, missing otherwise):

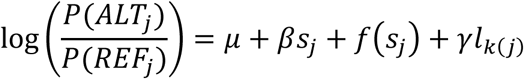

where *P*(*ALT*_*j*_) and *P*(*REF*_*j*_) are the fixation probabilities of the alternate (mutant) and reference (wild-type) alleles, respectively; *s*_*j*_ is the VEP score of variant *j*, β is its linear effect, *f*(*s*_*j*_) is a cubic spline function of the VEP score; *l*_*k*(*j*)_ is the local load (number of segregating singletons at chromosome arm *k* carrying variant *j*) as potential confounder, γ is its linear effect. Logistic GAMs were fit by the mgcv package in R.

## RESULTS

### Induced G:C-to-A:T point mutations were enriched for putative deleterious effects

Variants were classified as singletons if they passed our quality control and were observed in only one individual at the M_2_ generation. At the M_2_ generation, plants exposed to NaN_3_ had an average of 884 singletons, whereas the controls only had 12.5 singletons on average (Fig. 1). The average mutation rate in the SIEVE population is then estimated at 1 mutation per 308 kb, on par with reported mutation rates in a previous *B. distachyon* mutant population (24). As expected, the vast majority of singletons (94.2%) were G:C-to-A:T transitions (Fig. 2a), which are known to be preferentially induced by NaN_3_ (25, 37).

**Figure 1.**
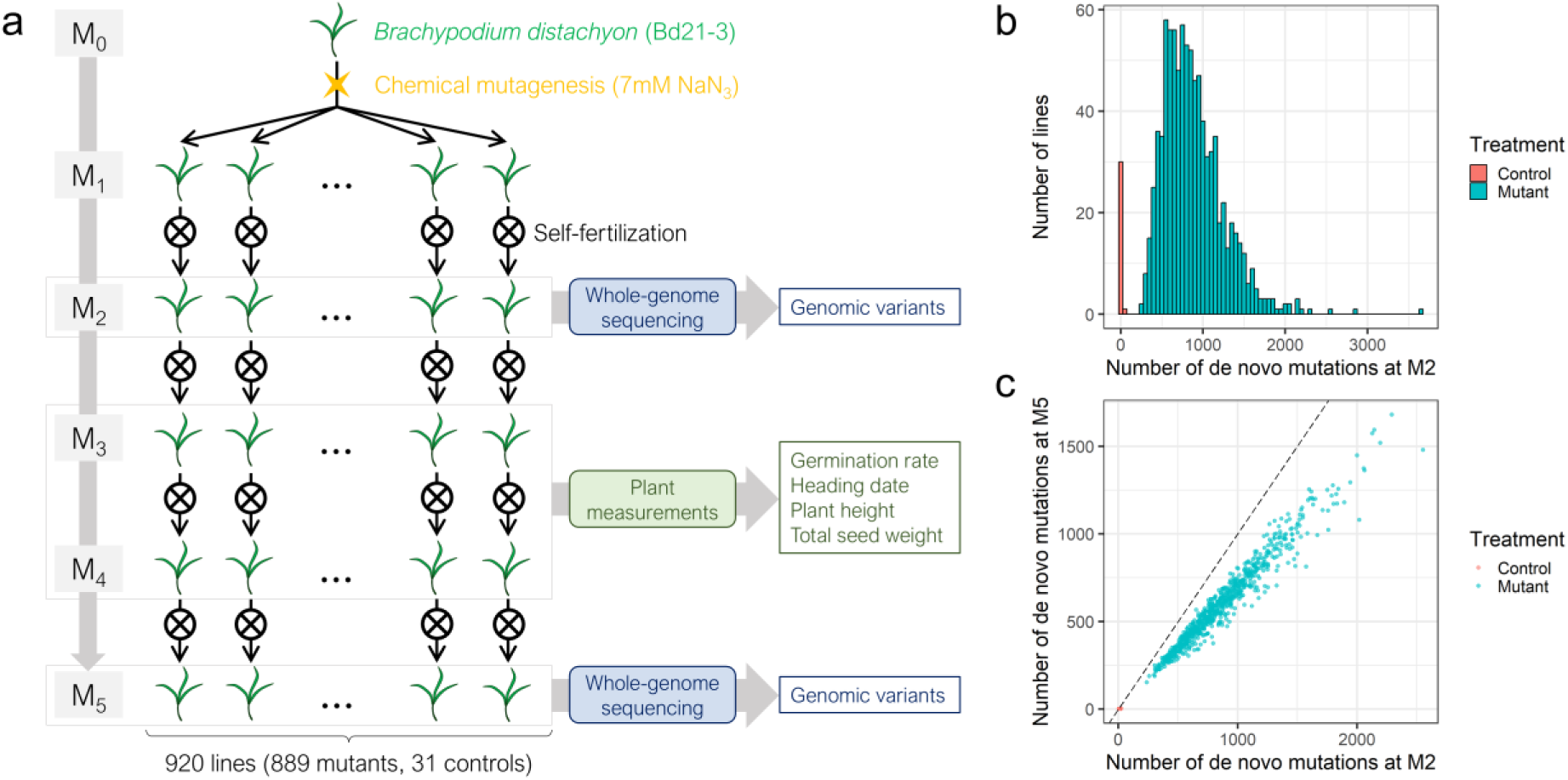
B*r*achypodium *distachyon* mutant population. **(a)** The *B. distachyon* mutant population (SIEVE) was developed by treating seeds with sodium azide (NaN_3_; 7 mM for mutants, 0 mM for controls) at the M_0_ generation, followed by 5 generations of selfing. Mutant and control lines were genotyped by whole-genome sequencing at generations M_2_ and M_5_ and phenotyped at the M_3_ and M_4_ generations. **(b)** Mutant lines carried on average 884 G:C-to-A:T singletons while control lines carried on average 12.5 G:C-to-A:T singletons. **(c)** The 858 lines (827 mutants, 31 controls) which survived to generation M_5_ carried on average 597.1 G:C-to-A:T singletons at M_5_ vs. 861.4 at M_2_ for mutants, and 1.5 G:C-to-A:T singletons at M_5_ vs. 12.5 at M_2_ for controls.

**Figure 2.**
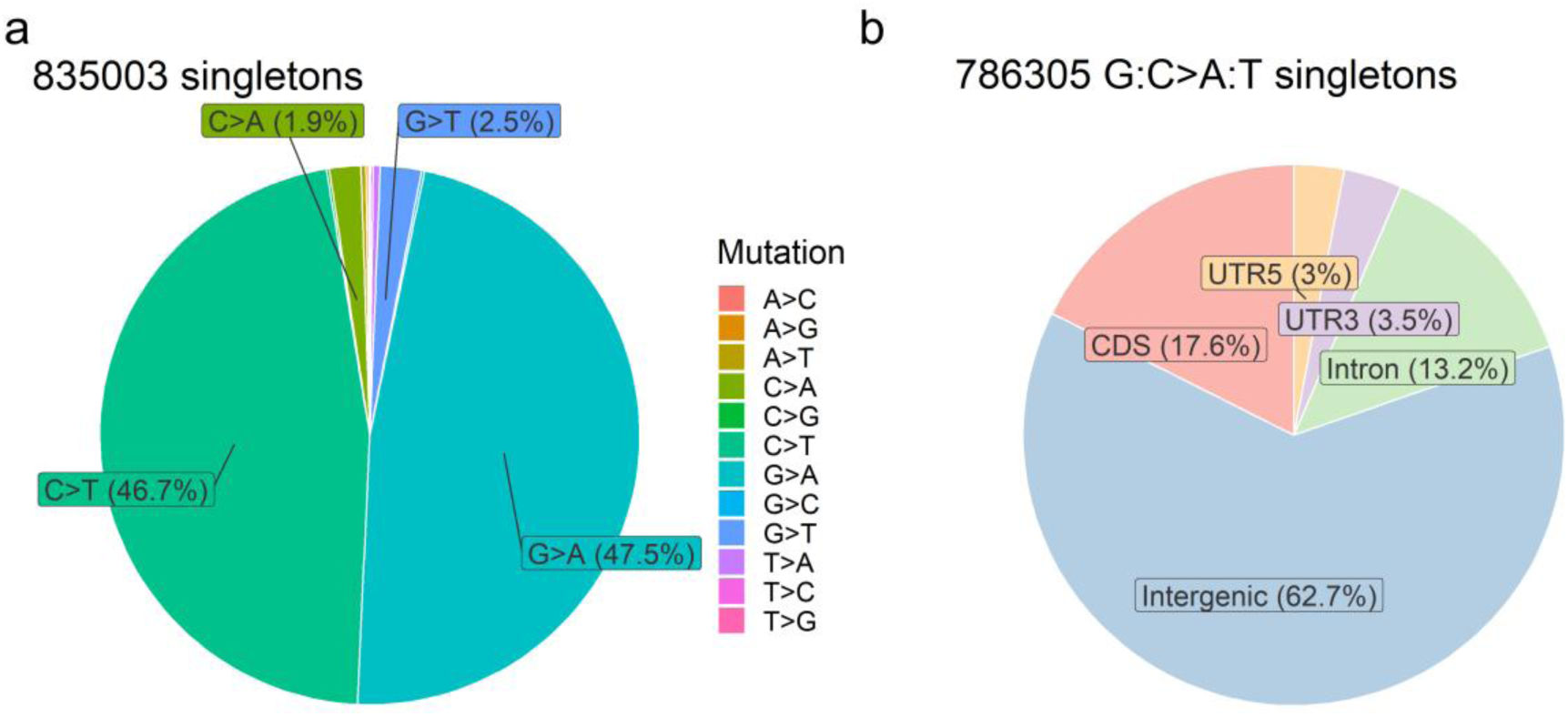
Classes of variants among singletons detected at the M_2_ generation. **(a)** Mutation type of observed singletons after quality control (genotype quality ≥ 20, read depth < 80). The vast majority (94.2%) of singletons were G:C-to-A:T transitions, as expected after mutagenesis by NaN_3_. **(b)** Gene features of G:C-to-A:T singletons. The majority (62.7%) of G:C-to-A:T singletons were in intergenic regions; CDS: coding sequence; UTR: untranslated region (3’ or 5’).

Hereafter, our analyses focused on the observed variants most likely to be induced by NaN_3_, i.e., G:C-to-A:T singletons. Among those mutations, the majority (62.7%) were in intergenic regions, while 17.6% were within protein-coding regions (Fig. 2b). Variant effects were predicted with different sequence-based VEP models depending on variant classes: missense, or ‘gene-proximal’, defined here as the non-coding variants flanking gene boundaries (4 kb upstream and 1 kb downstream of genes’ TSS, 1 kb upstream and 4 kb downstream of genes’ TTS) excluding protein-coding regions. As expected from theoretical and empirical distributions of fitness effects (38, 39), predicted fitness effects from SIFT and language models (ESM and PlantCAD) were skewed toward low values (Fig. 3). In contrast, predicted regulatory effects, from a2z and PhytoExpr (respectively for chromatin accessibility and RNA abundance), were distributed around zero with heavy tails (Fig. 3b). These distributions suggest that mutations in the SIEVE population may be deleterious, though only a small fraction of non-coding variants appear highly disruptive to regulatory processes.

**Figure 3.**
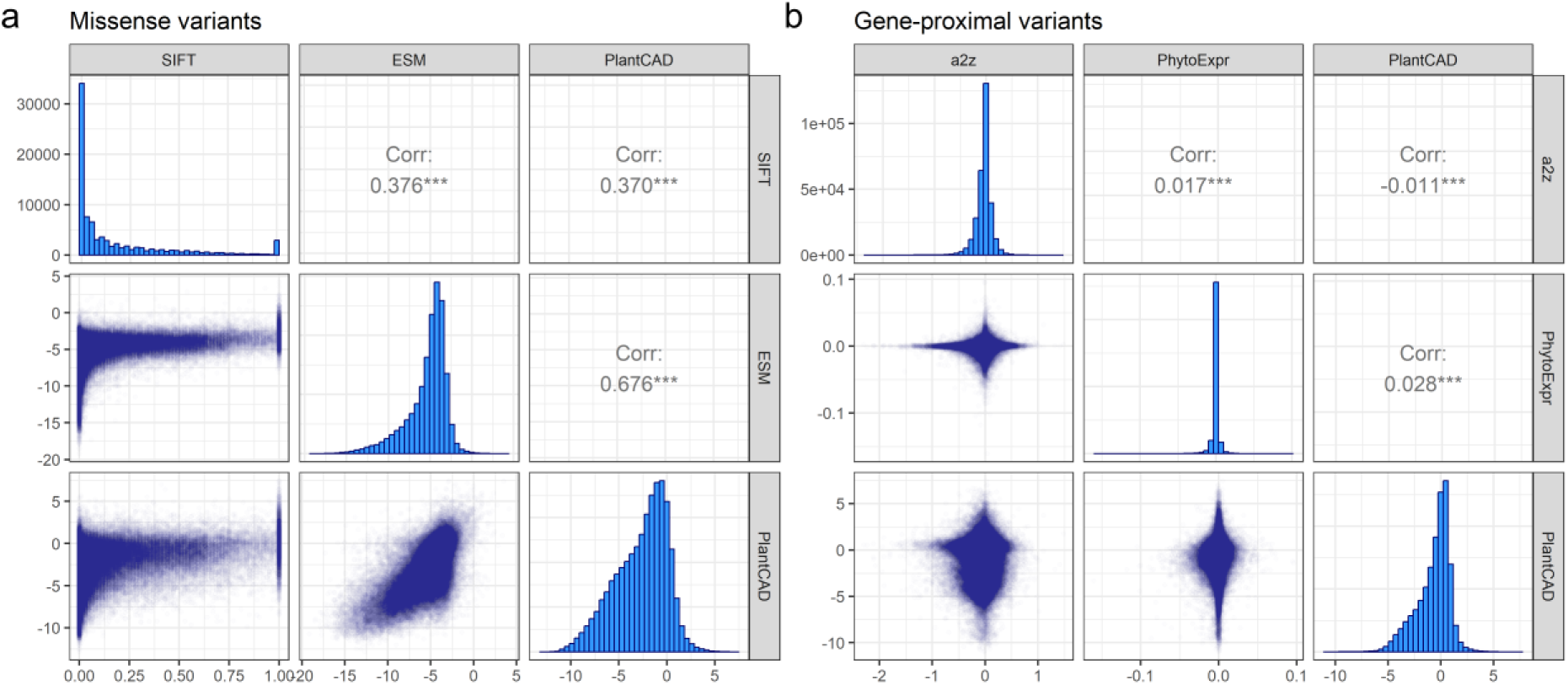
Distribution of predicted effects of G:C-to-A:T singletons at the M_2_ generation. **(a)** Predicted fitness effects of missense variants (variants causing amino-acid change in protein sequences of primary transcripts) from SIFT (multiple sequence alignments), ESM (protein language model), and PlantCAD (genomic language model). **(b)** Predicted effects of gene-proximal variants (variants within 4000 bp of a gene, excluding those in protein-coding regions) from a2z (chromatin-state model), PhytoExpr (sequence-to-expression model), and PlantCAD.

From generation M_2_ to M_5_, all 31 control lines survived, compared to 827 of the 889 mutant lines (93%). No significant difference in average VEP scores for missense variants was observed between surviving and lost lines (P > 0.32, *t*-test). Furthermore, only marginal differences in VEP scores for gene-proximal variants were detected for PlantCAD (-0.02 in surviving lines, P = 0.052) and a2z (+0.002, P = 0.026). Despite the large predicted effects of certain variants on fitness and regulation, genome-wide allele segregation from M_2_ to M_5_ among the 827 surviving mutant lines closely aligned with neutral expectations (Table 1).

**Table 1.** Allele segregation from M_2_ to M_5_.

| Genotype |  | Number of G:C-to-A:T singletons | Expected segregation from M<sub>2</sub> genotype (%) | Observed segregation from M<sub>2</sub> genotype (%) |
| --- | --- | --- | --- | --- |
| M<sub>2</sub> | M<sub>5</sub> |  |  |  |
| 0/1 | 0/0 | 218364 | 43.75 | 43.71 |
|  | 0/1 | 66046 | 12.5 | 13.22 |
|  | 1/1 | 215122 | 43.75 | 43.06 |
| 1/1 | 0/0 | 245 | 0 | 0.12 |
|  | 0/1 | 257 | 0 | 0.12 |
|  | 1/1 | 212349 | 100 | 99.76 |

0/0: homozygous wild-type genotype; 0/1: heterozygous genotype; 1/1: homozygous mutant genotype.
Percentages are based on 712,383 G:C-to-A:T singletons across 827 mutant lines.

This low level of segregation distortion suggests that selection on mutant alleles was relatively weak, likely due to small fitness effects of individual mutations or selective interference among multiple linked variants.

### Induced missense mutations were depleted in biological pathways related to central metabolism

Natural selection on the mutations observed at the M_2_ and M_5_ generations may provide information about fitness effects of genes. Specifically, here we determined if some classes of genes were less likely to carry induced missense mutations depending on their functional annotations. We considered GO annotations about biological processes, molecular functions, and cellular compartment, as well as PMN annotations about biological pathways. Gene depletion analyses at the M_2_ generation revealed that observed missense mutations were less likely in genes involved in central metabolic pathways: translation, DNA/protein metabolism, photosynthesis, and respiration (Table 2). Consistent with purifying selection acting through deleterious effects of mutations, depletions were generally stronger for homozygous mutations than heterozygous. Observed depletions in specific cellular compartments – ribosome, endoplasmic reticulum, thylakoid, and mitochondrion – were consistent with these pathways (Table S1). Gene depletion analyses for specific molecular functions revealed some negative selection in gene classes related to nucleotide binding. Interestingly, they also revealed relative enrichment of genes involved in ‘transcription regulator activity (GO:0140110)’ and ‘DNA-binding transcription factor activity (GO:0003700)’. These were the only functional classes in which gene enrichment was observed, indicating that mutations in transcription factor genes were under relatively weak selection.

**Table 2.** Gene depletion analysis of missense G:C-to-A:T singletons at generation M_2_: estimate and statistical significance of association between mutation density and gene class related to molecular pathways (GO BP or PMN annotations)

| Ontology | Gene class | Homozygous mutation density: coefficient (SE) | Heterozygous mutation density: coefficient (SE) | Likelihood ratio test: p-value (FDR) |
| --- | --- | --- | --- | --- |
| BP | translation (GO:0006412) | -95.3 (18.7) | -21.5 (9.2) | 2.9E-09 (9.5E-08) |
| BP | generation of precursor metabolites and energy (GO:0006091) | -66.3 (26.1) | -48 (15.6) | 6.4E-05 (0.0011) |
| BP | photosynthesis (GO:0015979) | -44.6 (27.4) | -64.5 (18.8) | 0.00017 (0.0018) |
| BP | metabolic process (GO:0008152) | -6.7 (3.9) | -5.7 (2.5) | 0.015 (0.08) |
| BP | protein metabolic process (GO:0019538) | -14.6 (6.1) | -6 (3.8) | 0.012 (0.08) |
| BP | DNA metabolic process (GO:0006259) | -14.6 (19) | -35.4 (13.3) | 0.014 (0.08) |
| PMN | NAD(P)/NADPH interconversion | -216.3 (86) | -97 (39.2) | 8.1E-05 (0.022) |
| PMN | aerobic respiration I (cytochrome c) | -65.2 (23.4) | -32.8 (13.3) | 0.00025 (0.033) |
| PMN | aerobic respiration III (alternative oxidase pathway) | -198.6 (77.3) | -68.6 (33.8) | 4E-04 (0.035) |
| PMN | ATP biosynthesis | -528.2 (222.7) | 19.7 (33.2) | 0.0015 (0.097) |

BP: Biological process (GO); PMN: Plant Metabolic Network (PlantCyc); SE: standard error; FDR: false discovery rate

We also sought to gain functional insights through allele purging from generations M_2_ to M_5_, to determine if missense variants in specific gene classes were under selection at later generations of population development. However, we detected no significant differences in purging probabilities across the investigated gene classes. This result may be due to small effects of missense variants in general, consistent with the low segregation distortion observed from M_2_ to M_5_ (Table 1). Additionally, the study may lack sufficient statistical power. A larger sample size or a higher mutation rate – resulting in a greater density of singletons in protein-coding regions – might have enabled the detection of purging differences with greater confidence.

### Putatively deleterious mutations negatively impacted whole-plant phenotypes at the M_3_ and M_4_ generations

To investigate the impact of induced mutations on whole-plant phenotypes, we recorded germination rate, plant height, total seed weight, and heading date during the M_3_ and M_4_ generations – stages at which plants were not disrupted by tissue sampling. Phenotypic correlations were generally low (-0.22 < R < 0.14 across generations), with the exception of plant height and total seed weight, which were strongly correlated (R = 0.65) (Figure S1). We subsequently assessed the association between these traits and mutation accumulation. The overall mutation load, defined as the total number of induced mutations per plant at M_2_, correlated strongly with deleterious effects on recorded phenotypes (Table 3). Specifically, increased mutation load was associated with reductions in germination rate, plant height, and total seed weight, as well as delayed heading dates. As expected, the phenotypic impact of the homozygous load was substantially greater than that of the heterozygous load, except for germination rate for which effects of both loads were similar.

**Table 3.**
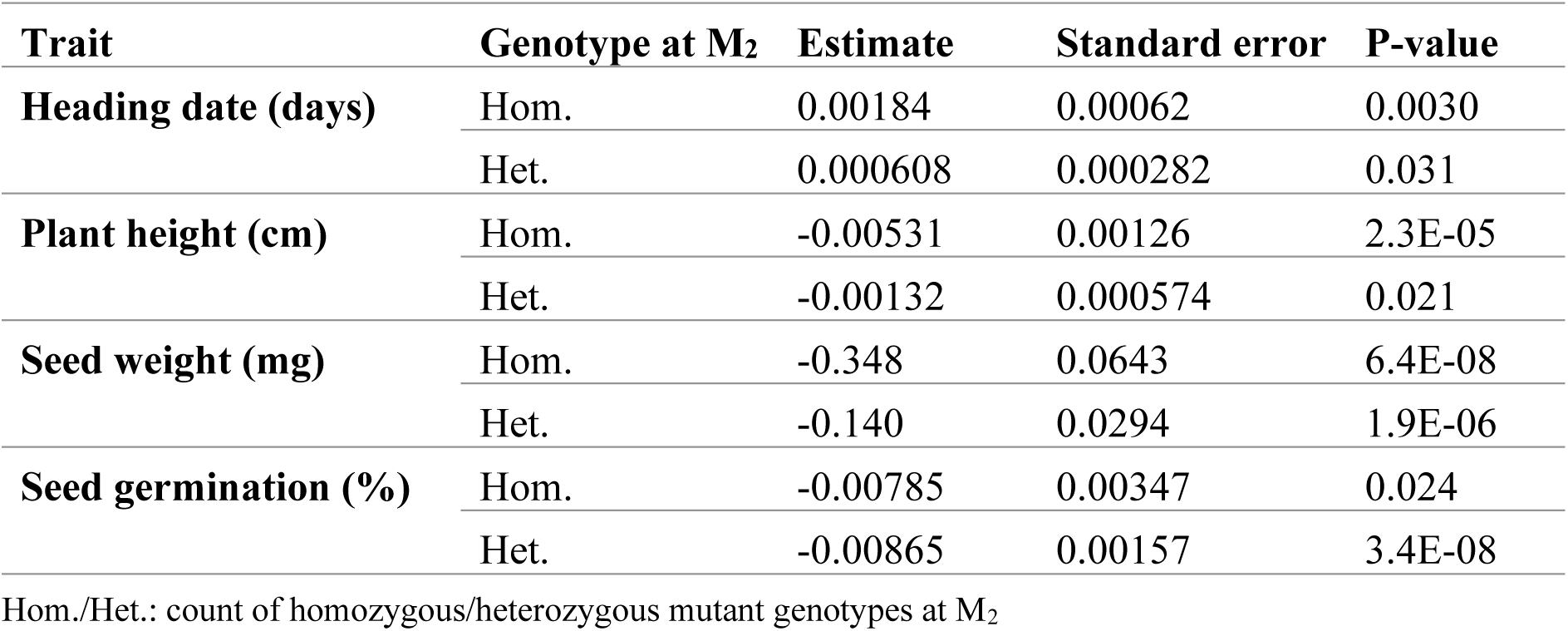
Effect of mutational burden on phenotypic traits at generations M_3_ and M_4_.

To determine if a subset of high-impact variants exerted a disproportionate influence on whole-plant phenotypes, we modeled mutation burden across varying VEP score thresholds while controlling for background mutation effects by including total mutation load as a covariate. These models revealed that the phenotypic impact of homozygous mutations intensified with increasing threshold stringency, particularly for variants with highly negative VEP scores (Fig. 4).

**Figure 4.**
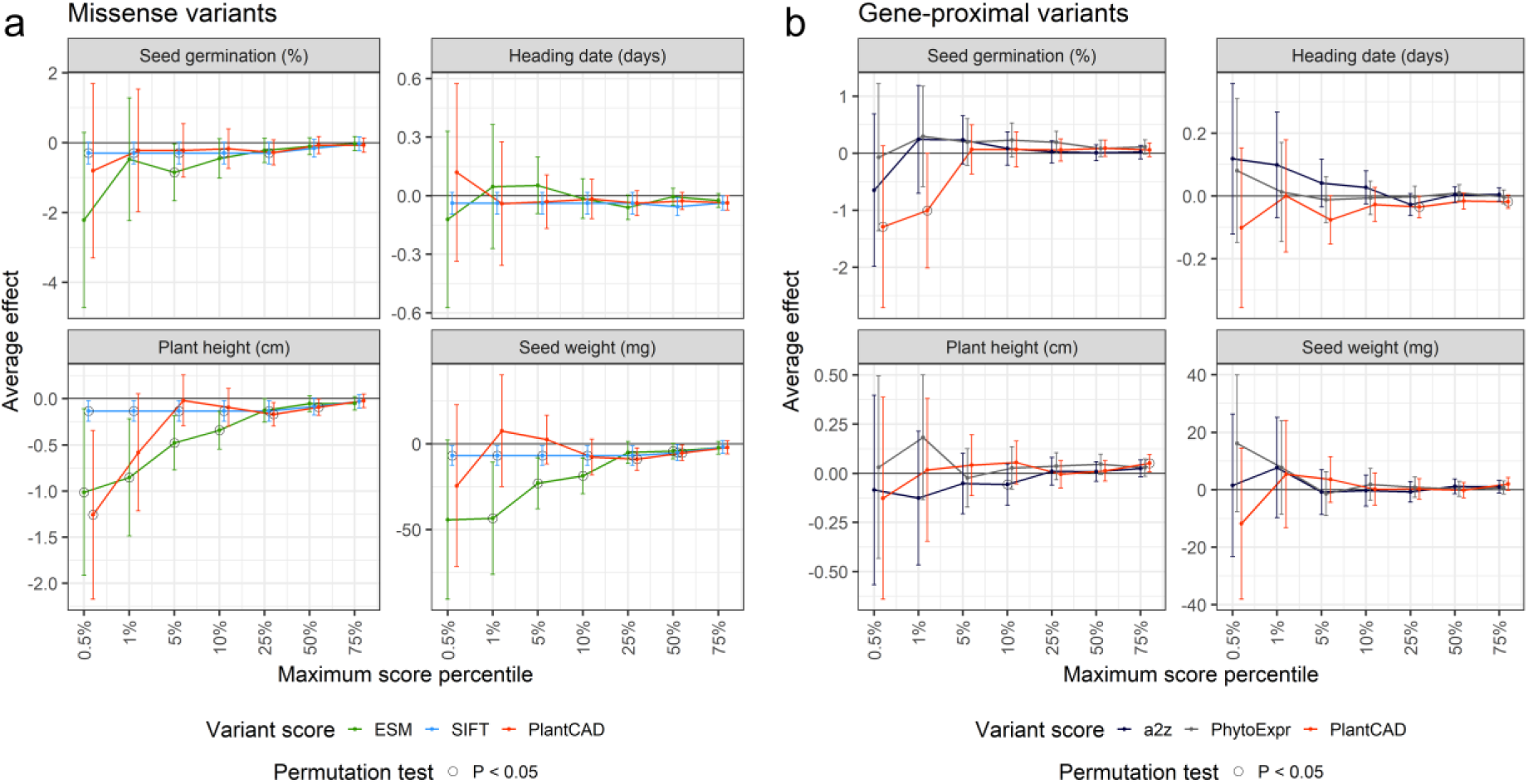
Functional enrichment of homozygous variants prioritized for predicted negative effects. Average effect (point estimate + 95% confidence interval) of homozygous G:C-to-A:T singletons on seed germination, heading date, plant height, and total seed weight at the M_3_ and M_4_ generations. Black circles refer to average effects that are significant based on 100 permutations of predicted variant effects (empirical P-value lower than 0.05). **(a)** Prioritizations of missense variants (variants causing amino-acid change in protein sequences of primary transcripts) by SIFT (multiple sequence alignments), ESM (protein language model), or PlantCAD (genomic language model). **(b)** Prioritization of gene-proximal variants (variants within 4000 bp of a gene, excluding those in protein-coding regions) by a2z (chromatin-state model), PhytoExpr (sequence-to-expression model), or PlantCAD.

Consistent with the expected correlation between predicted scores and fitness, the average effect of missense mutations within the bottom 5^th^ percentile (thresholds: ESM = -10.3, SIFT = 0, PlantCAD = -7.8) was significantly negative for germination rate (SIFT, ESM), plant height (SIFT, ESM, PlantCAD), and total seed weight (SIFT, ESM) (Fig. 4a). These results suggest that scores from ESM, a protein language model, were particularly effective at capturing the deleterious effects of homozygous missense variants, outperforming both SIFT (an MSA-based approach) and PlantCAD (a genomic language model).

Among gene-proximal variants, the average effect of variants within the bottom 5^th^ percentile of VEP scores (thresholds: PlantCAD = -3.9, a2z = -0.3, PhytoExpr = -0.006) generally did not deviate significantly from zero. A notable exception was observed for variants prioritized by PlantCAD, which exhibited a significantly negative effect on germination rate (Fig. 4b).

When prioritizing variants with highly positive VEP scores, significant phenotypic effects were detected only among gene-proximal variants (Fig. S2). Unexpectedly, variants within the top 95th percentile of PlantCAD scores (threshold: 1.2) were associated with reduced plant height, decreased seed weight, and delayed heading dates (Fig. S2b). This result is counterintuitive, as positive PlantCAD scores are intended to identify variants with beneficial effects.

Altogether, our burden tests indicate that prioritizing variants with low VEP scores – particularly those in the bottom 5^th^ percentile of ESM scores – effectively captures deleterious effects on whole-plant phenotypes. However, our results also suggest that prioritizing variants with high VEP scores (e.g., PlantCAD scores above the 95^th^ percentile) does not reliably identify beneficial phenotypic effects. It should be noted, however, that our dataset may be underpowered to detect beneficial variant effects, as the distribution of VEP scores in the SIEVE population is heavily skewed toward deleterious values (Fig. 3).

### Predicted effects of mutations were associated with their observed fitness from generations M_2_ to M_5_

While our burden tests evaluated the capacity of predicted variant effects to capture phenotypic variation, they did not establish a direct functional relationship between VEP scores and evolutionary fitness. To address this, we conducted independent purging tests by fitting VEP scores to the fixation probability of mutant alleles from M_2_ to M_5_ Specifically, we modeled the ’relative fixation probability,’ defined as the log-ratio of the fixation probability of the mutant allele over that of the wild type. Theoretically, the relative fixation probability for a given variant *j* equals log (*w*_*j*_), where *w*_*j*_ is the fitness of the mutant allele relative to the wild type. Consequently, these analyses allow us to map VEP scores directly to the relative fitness of variants within our experimental population.

Among missense variants, the association between relative fixation probability and VEP scores was significant for both ESM and PlantCAD (P < 0.05, likelihood ratio test; Fig. 5a). The stronger signal observed for ESM scores (P = 0.005 vs. P = 0.036 for PlantCAD) is consistent with our phenotypic burden tests (Fig. 4a), further suggesting that ESM is more effective at capturing the fitness consequences of missense mutations. Interestingly, the inferred relationships between VEP scores and the relative fixation probability of missense variants did not deviate significantly from linearity (P > 0.16, Wald test; Table 4). This suggests a log-linear relationship between the VEP score of a missense variant *j* and its relative fitness *w*_*j*_. This finding is especially meaningful for ESM and PlantCAD, as these scores correspond to a log-odds ratio 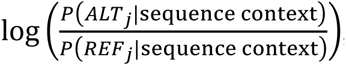, mirroring the mathematical structure of the relative fixation probability.

**Figure 5.**
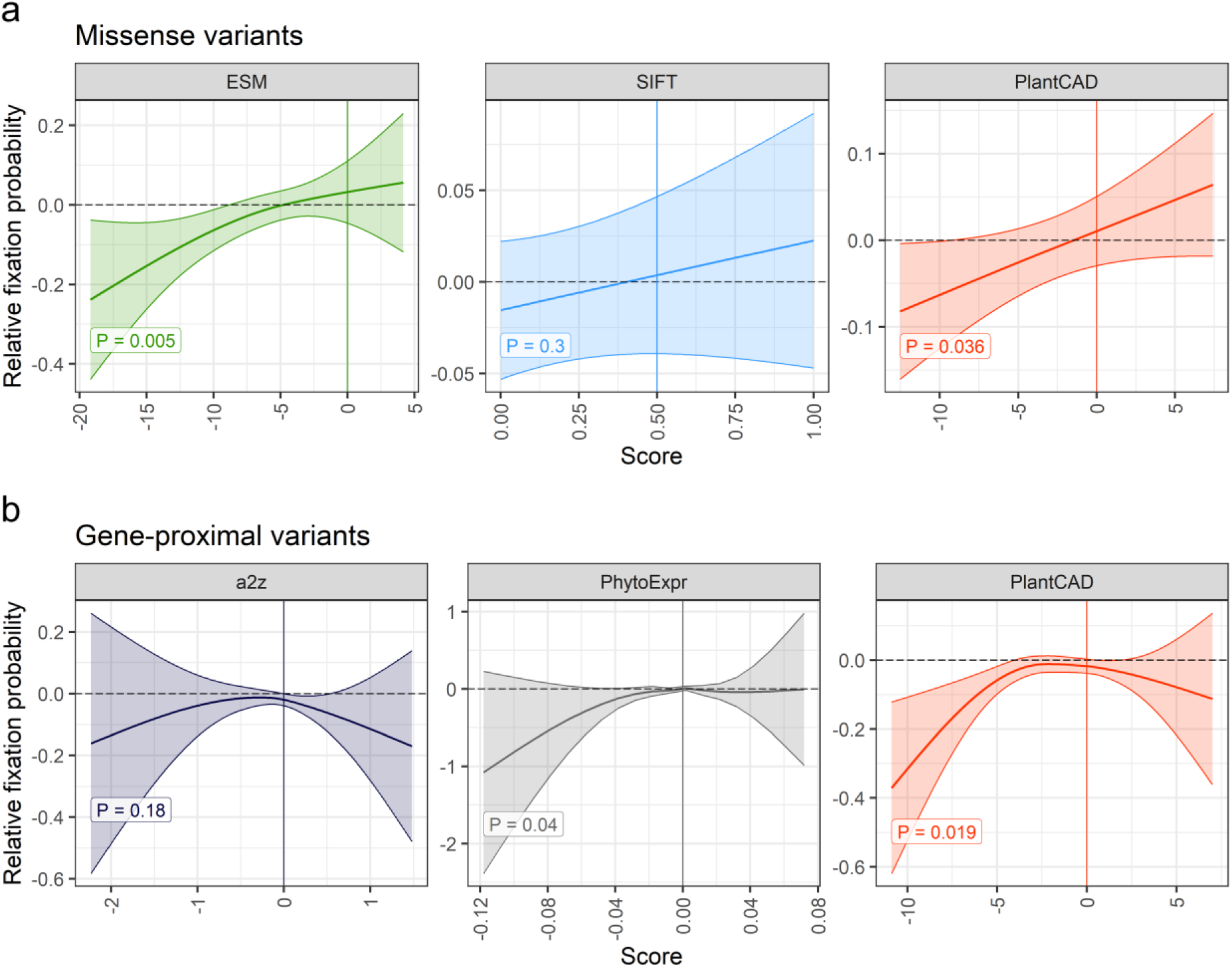
Association between observed fitness of variant and predicted variant effect. Relative fixation probability: 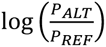, where *P_ALT_* is the probability that a heterozygous variant at M_2_ generation is homozygous for the alternate (mutant) allele at M_5_ generation; similarly for *P*_*REF*_ and the reference (wild-type) allele. P: P-value from significance test on the relationship between relative fixation probability and variant scores, as inferred from a logistic generalized additive model. **(a)** Scores of missense variants (variants causing amino-acid change in protein sequences of primary transcripts by SIFT (multiple sequence alignments), ESM (protein language model), or PlantCAD (genomic language model). **(b)** Scores of gene-proximal variants (variants within 4000 bp of a gene, excluding those in protein-coding regions) by a2z (chromatin-state model), PhytoExpr (sequence-to-expression model), or PlantCAD.

**Table 4.**
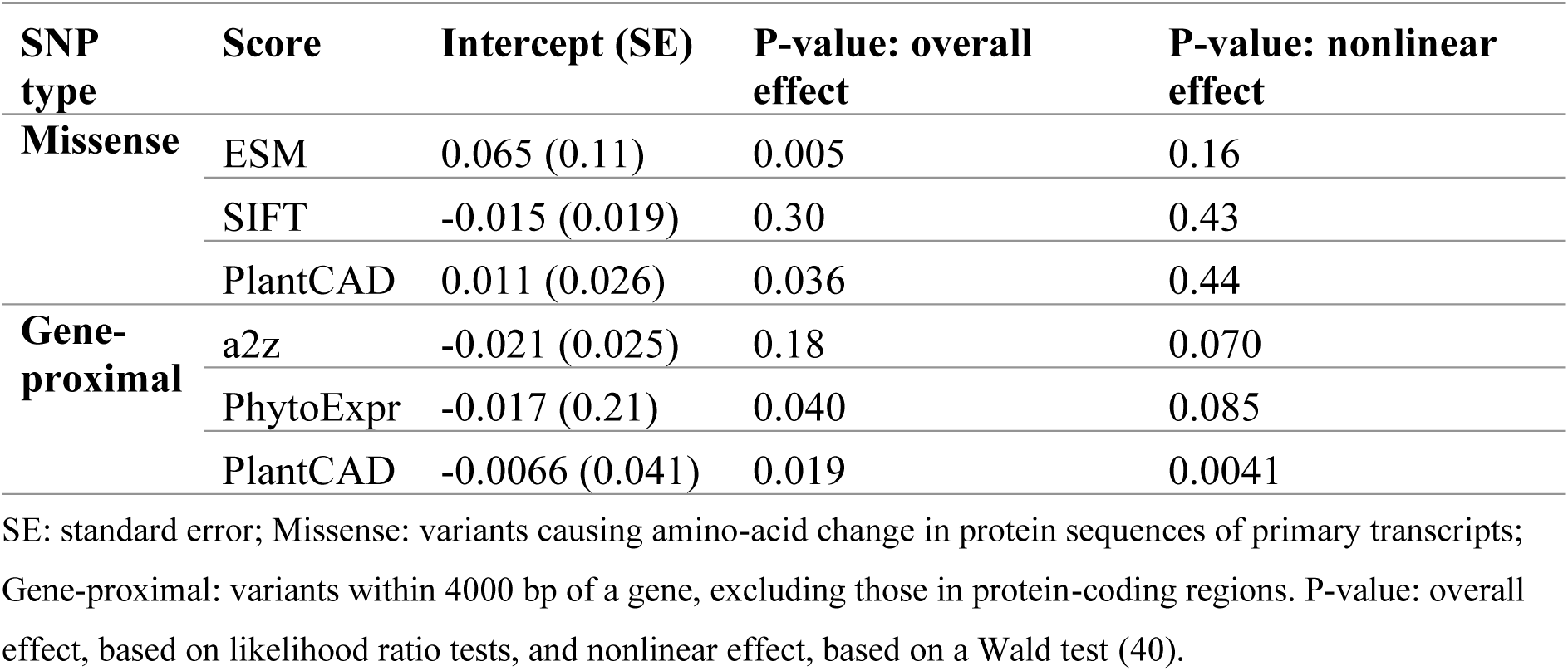
Generalized Additive Model (GAM) fit for fixation probability of mutant allele depending on variant effect prediction.

Among gene-proximal variants, the association between relative fixation probability and VEP scores was non-significant for the chromatin-state model a2z. While the association was significant for the sequence-to-expression model PhytoExpr, the signal was weak, with the confidence interval encompassing zero across the range of scores (Fig. 5b). In contrast, PlantCAD showed a significant and relatively strong association. However, unlike the trend observed for missense variants, this relationship deviated significantly from linearity (P = 0.0041, Wald test; Table 4). Notably, PlantCAD scores above -5 failed to correlate with relative fixation probability, indicating a limited capacity to distinguish neutral from beneficial variants in that range. This non-linearity may be driven by the functional heterogeneity of gene-proximal variants, which can influence gene activity through diverse biological mechanisms (e.g., enhancer, promoter, initiation, termination, or splicing activity), unlike the more uniform impact of missense mutations.

Remarkably, for all VEP scores and variant sets analyzed, the predicted relative fixation probability did not differ significantly from zero when the score was null (0.5 for SIFT; 0 for others). This indicates that all VEP models are well-calibrated, in that a score of zero (or 0.5 for SIFT) accurately reflects neutral fitness effects.

Collectively, our burden and purging tests suggest that ESM scores are the most robust predictors of variant effects on evolutionary fitness and fitness-related traits (e.g., total seed weight) among missense variants. While PlantCAD remains a promising tool, the counterintuitive results associated with positive values in both burden and purging tests suggest that further validation is required to assess its ability to reliably detect beneficial effects.

## DISCUSSION

### Validating predicted variant effects in mutant populations

In this study, we developed the SIEVE population specifically to evaluate variant effect predictions at base-pair resolution. To this end, we proposed different statistical analyses to validate the impact of candidate variants: (*i*) gene depletion analyses to identify gene classes under selection; (*ii*) burden tests to quantify the association between mutation loads (prioritized by VEP) and phenotypes; and (*iii*) purging tests to establish the relationship between VEP scores and the relative fixation probability of mutant alleles.

While previous studies in crops such as maize, potato, and cassava have successfully detected associations between predicted variant effects and field traits (20, 41–43), those investigations relied on natural accessions. In such diverse populations, extensive physical linkage – spanning more than 10 kb – often confounds the resolution of variant-trait associations, making it difficult to demonstrate VEP accuracy at the level of individual variants. Furthermore, accessions in these studies are typically genotyped by mapping reads to a single reference genome. This approach introduces significant reference bias, as loci unique to diverse genomic backgrounds are omitted if they are absent from the reference assembly (44, 45). The SIEVE population addressed both of these limitations (LD-block resolution and reference bias) by generating genetically independent mutant lines derived from a single, high-quality reference genotype (Bd21-3). Unlike existing mutant resources in species like Brachypodium, barley, wheat, or maize (24, 26, 46, 47), the genomic and phenotypic data acquired across generations in SIEVE provided the necessary resources for validation and calibration of VEP models. Specifically, our data enabled the implementation of burden and purging tests, which allowed us to compare predicted variant effects with realized impacts on plant fitness.

Crucially, the deep-learning VEP models evaluated here were trained in multiple species across the tree of life (ESM) or the angiosperm clade (a2z, PhytoExpr, PlantCAD), rather than specifically on Brachypodium (5, 10–12). Consequently, the successful validation of these models in the SIEVE population should support their broad application across related taxa, particularly in crops closely related to Brachypodium, like wheat, barley, and rice.

Our gene depletion analyses at the M_2_ generation proved effective to identify genes involved in essential functions. However, we did not observe significant deviations in fixation probabilities across gene classes from M_2_ to M_5_. This lack of signal likely stems from insufficient statistical power to detect small-effect variants within specific gene categories. Future efforts aiming to resolve this issue will require even larger populations of mutant lines. Additionally, the observed lack of selection pressure on specific gene classes may be attributed to our focus on fitness under non-stressful growth conditions (48). Applying specific biotic or abiotic stressors (e.g., drought, pathogen pressure) in future studies could amplify the selection pressure on genes of interest and facilitate the identification of variants involved in targeted stress-tolerance traits.

### Predicting fitness effects of variants with biological language models

Consistent with previous benchmarks of variant pathogenicity in humans (6, 49), biological LMs proved particularly accurate for predicting variant effects in this study. Among these, the protein LM ESM demonstrated higher predictive accuracy than the genomic LM PlantCAD in both burden and purging tests for missense variants. Consequently, according to our validations, protein LMs currently represent the most promising tool for VEP, with outstanding potential in research and breeding applications. Nevertheless, genomic LMs offer distinct advantages that make them particularly attractive for VEP.

First, genomic LMs are inherently more versatile, as they can predict the effects of non-coding mutations. In our study, low PlantCAD scores for gene-proximal variants were significantly associated with negative impacts on both germination rate (Fig. 4b) and overall fitness (Fig. 5b), highlighting the model’s utility for identifying deleterious variants outside of protein-coding regions. Second, their predictions are independent of transcript annotation quality. The reliance of protein LMs on accurate annotations can be a significant hurdle when working with *de novo* genome assemblies or non-model species where high-quality transcriptomes are unavailable. As *de novo* assemblies become increasingly common in plant genomics (50), ‘annotation-free’ VEP techniques like PlantCAD will likely become essential.

Despite key advantages, genomic LMs still face specific challenges, particularly when applied to functionally heterogeneous variant sets in non-coding regions. Both burden and purging tests revealed that positive PlantCAD scores of gene-proximal variants did not correlate with beneficial fitness effects. Future research should investigate whether data curation practices during pre-training influenced this lack of directionality. For example, genomic LMs such as GPN and PlantCAD are often pre-trained on datasets where non-coding regions and repetitive sequences are down-sampled or down-weighted (1, 10, 21). While these practices are intended to mitigate genomic biases, their specific impact on VEP accuracy in non-coding regions has yet to be established.

#### Implementing variant effect prediction in precision breeding

Our aim in this study was to evaluate VEP models for their deployment in crop improvement. The successful validation of models like ESM should motivate its application for both genomic prediction and genome editing.

In genomic prediction, modeling the load of deleterious mutations is a promising strategy to select individuals with enhanced fitness (20, 41, 51). Future prediction models could leverage our insight into the relationship between VEP scores and fitness-related traits to estimate the effect of variants on key breeding traits. Our burden tests indicated that missense variants with an ESM score below the 10^th^ percentile (threshold: -8.7) exerted a significantly more deleterious impact on plant height and total seed weight (Fig. 4). This threshold aligns with the value of -7 proposed previously (6, 52), which could serve as a standard criterion for identifying a subset of putatively deleterious variants to be upweighted in genomics-assisted selection. Alternatively, breeders might infer a mathematical function to relate VEP scores to variant effects on fitness-related traits. Our purging tests suggest that the relationship between missense VEP scores (e.g., ESM or PlantCAD) and fitness is best modeled on a log-scale, as evidenced by the linear relationship between these scores and the logarithm of allele fitness. Our results provide a hypothesis about the relationship between VEP scores and variant effects, which remains to be tested in further studies on realized phenotypic effects under field conditions.

Beyond genomic prediction, VEP scores may also be used to prioritize specific variants for targeted mutagenesis and genome editing. Our findings demonstrate that state-of-the-art VEP models can detect genetic effects at the resolution of individual mutations, suggesting they could be applied for targeting beneficial alleles. In such applications, breeders may prioritize ‘back mutations’, that is, variants reverting derived, deleterious alleles back to their ancestral state (53). Natural candidates for this ‘targeted purging’ of deleterious variants are DNA transitions (G:C-to-A:T or A:T-to-G:C), for two primary reasons: first, G:C-to-A:T mutations were the predominant class of variants induced in this study (Fig. 2); and second, cytidine (C-to-T) and adenosine (A-to-G) base editors are currently the most advanced base-editing tools in crops (54, 55). Notably, recent work has successfully used SIFT scores to target and fix a deleterious variant in tomato (56). Proof-of-concept studies leveraging more advanced VEP models, such as ESM or PlantCAD, will be essential to determine the efficiency of *in silico* variant prioritization for targeted purging. In particular, future studies in diverse crops are needed to estimate the proportion of variants leading to detectable impacts on traits of interest, among those prioritized by sequence-based deep learning techniques.

Our study focused on fitness-related traits, which certainly explains why sequence-to-function models, such as a2z and PhytoExpr, were less effective than biological LMs at capturing the phenotypic impacts observed in the SIEVE population (1). However, these models remain valuable to investigate specific genes of interest, particularly when identifying candidate variants for targeted follow-up analyses. For example, sequence-to-function models could be applied to predict the impact of specific up-regulating variants on the function of a target gene. In such research contexts, the SIEVE population may serve as a valuable genetic resource for future candidate variant analyses (57).

## Acknowledgements

This research is supported by the Novo Nordisk Foundation, Grant NNF21OC0067311.

## Data availability statement

The input datasets used in this study are publicly available on Zenodo (58). The scripts used to conduct analyses (genotype calling, variant scoring, mutant analyses) are available on GitHub (https://github.com/guillaume-ramstein/sieve/tree/main).

## References

1. J. Sendrowski, T. Bataillon, G. P. Ramstein, In silico prediction of variant effects: promises and limitations for precision plant breeding. Züchter Genet. Breed. Res. 138, 1–20 (2025).

2. Z. Joly-Lopez, J. M. Flowers, M. D. Purugganan, Developing maps of fitness consequences for plant genomes. Curr. Opin. Plant Biol. 30, 101–107 (2016).

3. A. Rives, et al., Biological structure and function emerge from scaling unsupervised learning to 250 million protein sequences. Proc. Natl. Acad. Sci. U. S. A. 118 (2021).

4. G. Benegas, C. Ye, C. Albors, J. C. Li, Y. S. Song, Genomic language models: opportunities and challenges. Trends Genet. 41, 286–302 (2025).

5. J. Meier, et al., Language models enable zero-shot prediction of the effects of mutations on protein function. bioRxiv 34, 29287–29303 (2021).

6. N. Brandes, G. Goldman, C. H. Wang, C. J. Ye, V. Ntranos, Genome-wide prediction of disease variant effects with a deep protein language model. Nat. Genet. 55, 1512–1522 (2023).

7. Y. Bromberg, R. Prabakaran, A. Kabir, A. Shehu, Variant effect prediction in the age of machine learning. Cold Spring Harb. Perspect. Biol. 16, a041467 (2024).

8. D. R. Tabet, et al., Benchmarking computational variant effect predictors by their ability to infer human traits. Genome Biol. 25, 172 (2024).

9. R. Vaser, S. Adusumalli, S. N. Leng, M. Sikic, P. C. Ng, SIFT missense predictions for genomes. Nat. Protoc. 11, 1–9 (2016).

10. J. Zhai, et al., Cross-species modeling of plant genomes at single-nucleotide resolution using a pretrained DNA language model. Proc. Natl. Acad. Sci. U. S. A. 122, e2421738122 (2025).

11. T. Wrightsman, A. P. Marand, P. A. Crisp, N. M. Springer, E. S. Buckler, Modeling chromatin state from sequence across angiosperms using recurrent convolutional neural networks. Plant Genome 15, e20249 (2022).

12. T. Li, et al., Modeling 0.6 million genes for the rational design of functional cis-regulatory variants and de novo design of cis-regulatory sequences. Proc. Natl. Acad. Sci. U. S. A. 121, e2319811121 (2024).

13. A. R. Mansisidor, V. I. Risca, Chromatin accessibility: methods, mechanisms, and biological insights. Nucleus 13, 236–276 (2022).

14. M. E. Hauberg, et al., Common schizophrenia risk variants are enriched in open chromatin regions of human glutamatergic neurons. Nat. Commun. 11, 5581 (2020).

15. Z. Lu, et al., The prevalence, evolution and chromatin signatures of plant regulatory elements. Nat Plants 5, 1250–1259 (2019).

16. E. Rodgers-Melnick, D. L. Vera, H. W. Bass, E. S. Buckler, Open chromatin reveals the functional maize genome. Proc. Natl. Acad. Sci. U. S. A. 113, E3177–84 (2016).

17. A. Patel, et al., DART-Eval: A comprehensive DNA language model evaluation benchmark on regulatory DNA. arXiv [cs.LG] (2024).

18. J. D. Washburn, et al., Evolutionarily informed deep learning methods for predicting relative transcript abundance from DNA sequence. Proc. Natl. Acad. Sci. U. S. A. 116, 5542–5549 (2019).

19. F. F. Peleke, S. M. Zumkeller, M. Gültas, A. Schmitt, J. Szymański, Deep learning the cis-regulatory code for gene expression in selected model plants. Nat. Commun. 15, 3488 (2024).

20. G. P. Ramstein, E. S. Buckler, Prediction of evolutionary constraint by genomic annotations improves functional prioritization of genomic variants in maize. Genome Biol. 23, 1–26 (2022).

21. G. Benegas, S. S. Batra, Y. S. Song, DNA language models are powerful predictors of genome-wide variant effects. Proc. Natl. Acad. Sci. U. S. A. 120, e2311219120 (2023).

22. B. S. Gaut, A. D. Long, The lowdown on linkage disequilibrium. Plant Cell 15, 1502–1506 (2003).

23. S. A. Flint-Garcia, J. M. Thornsberry, E. S. Buckler 4th, Structure of linkage disequilibrium in plants. Annu. Rev. Plant Biol. 54, 357–374 (2003).

24. M. Dalmais, et al., A TILLING Platform for Functional Genomics in Brachypodium distachyon. PLoS One 8, e65503 (2013).

25. I. Szarejko, et al., “Creation of a TILLING population in barley after chemical mutagenesis with sodium azide and MNU” in Biotechnologies for Plant Mutation Breeding, (Springer, Cham, 2017), pp. 91–111.

26. J. Zhang, et al., Sequencing 4.3 million mutations in wheat promoters to understand and modify gene expression. Proc. Natl. Acad. Sci. U. S. A. 120, e2306494120 (2023).

27. J. Brkljacic, et al., Brachypodium as a model for the grasses: today and the future. Plant Physiol. 157, 3–13 (2011).

28. M. W. Bevan, D. F. Garvin, J. P. Vogel, Brachypodium distachyon genomics for sustainable food and fuel production. Curr. Opin. Biotechnol. 21, 211–217 (2010).

29. J. C. Zadoks, T. T. Chang, C. F. Konzak, A decimal code for the growth stages of cereals. Weed Res. 14, 415–421 (1974).

30. H. Li, Aligning sequence reads, clone sequences and assembly contigs with BWA-MEM. arXiv [q-bio.GN] (2013).

31. A. McKenna, et al., The Genome Analysis Toolkit: a MapReduce framework for analyzing next-generation DNA sequencing data. Genome Res. 20, 1297–1303 (2010).

32. X. Zheng, et al., A high-performance computing toolset for relatedness and principal component analysis of SNP data. Bioinformatics 28, 3326–3328 (2012).

33. X. Zheng, B. S. Weir, Eigenanalysis of SNP data with an identity by descent interpretation. Theor. Popul. Biol. 107, 65–76 (2016).

34. P. C. Ng, S. Henikoff, Predicting deleterious amino acid substitutions. Genome Res. 11, 863–874 (2001).

35. C. Hawkins, et al., Plant Metabolic Network 15: A resource of genome-wide metabolism databases for 126 plants and algae. J. Integr. Plant Biol. 63, 1888–1905 (2021).

36. F. Laporte, A. Charcosset, T. Mary-Huard, Efficient ReML inference in variance component mixed models using a Min-Max algorithm. PLoS Comput. Biol. 18, e1009659 (2022).

37. O. Olsen, X. Wang, D. von Wettstein, Sodium azide mutagenesis: preferential generation of A.T-->G.C transitions in the barley Ant18 gene. Proc. Natl. Acad. Sci. U. S. A. 90, 8043–8047 (1993).

38. C. Bank, G. B. Ewing, A. Ferrer-Admettla, M. Foll, J. D. Jensen, Thinking too positive? Revisiting current methods of population genetic selection inference. Trends Genet. 30, 540–546 (2014).

39. J. Chen, T. Bataillon, S. Glémin, M. Lascoux, What does the distribution of fitness effects of new mutations reflect? Insights from plants. New Phytol. 233, 1613–1619 (2022).

40. S. N. Wood, On p-values for smooth components of an extended generalized additive model. Biometrika 100, 221–228 (2013).

41. J. Yang, et al., Incomplete dominance of deleterious alleles contributes substantially to trait variation and heterosis in maize. PLoS Genet. 13, e1007019 (2017).

42. Y. Wu, et al., Phylogenomic discovery of deleterious mutations facilitates hybrid potato breeding. Cell 186, 2313–2328.e15 (2023).

43. E. M. Long, M. C. Romay, G. Ramstein, E. S. Buckler, K. R. Robbins, Utilizing evolutionary conservation to detect deleterious mutations and improve genomic prediction in cassava. Front. Plant Sci. 13 (2023).

44. J. L. Gage, et al., Multiple maize reference genomes impact the identification of variants by genome-wide association study in a diverse inbred panel. Plant Genome 12, 180069 (2019).

45. R. Della Coletta, Y. Qiu, S. Ou, M. B. Hufford, C. N. Hirsch, How the pan-genome is changing crop genomics and improvement. Genome Biol. 22, 3 (2021).

46. M. Schreiber, et al., A highly mutagenised barley (cv. Golden Promise) TILLING population coupled with strategies for screening-by-sequencing. Plant Methods 15, 99 (2019).

47. Y. Wang, et al., Large DNA and protein language models enhance discovery of deleterious mutations in maize. Genome Biol. 26, 412 (2025).

48. A. F. Agrawal, M. C. Whitlock, Environmental duress and epistasis: how does stress affect the strength of selection on new mutations? Trends Ecol. Evol. 25, 450–458 (2010).

49. G. Benegas, C. Albors, A. J. Aw, C. Ye, Y. S. Song, A DNA language model based on multispecies alignment predicts the effects of genome-wide variants. Nat. Biotechnol. 1–6 (2025). 10.1038/s41587-024-02511-w.

50. M. Schreiber, M. Jayakodi, N. Stein, M. Mascher, Plant pangenomes for crop improvement, biodiversity and evolution. Nat. Rev. Genet. (2024). 10.1038/s41576-024-00691-4.

51. T. J. Y. Kono, et al., The Role of Deleterious Substitutions in Crop Genomes. Mol. Biol. Evol. 33, 2307–2317 (2016).

52. C. M. Andorf, et al., PanEffect: a pan-genome visualization tool for variant effects in maize. Bioinformatics 40 (2024).

53. T. Latrille, J. Joseph, D. A. Hartasánchez, N. Salamin, Estimating the proportion of beneficial mutations that are not adaptive in mammals. PLoS Genet. 20, e1011536 (2024).

54. H. Zhu, C. Li, C. Gao, Applications of CRISPR-Cas in agriculture and plant biotechnology. Nat. Rev. Mol. Cell Biol. (2020). 10.1038/s41580-020-00288-9.

55. B. Li, C. Sun, J. Li, C. Gao, Targeted genome-modification tools and their advanced applications in crop breeding. Nat. Rev. Genet. 25, 603–622 (2024).

56. A. N. Glaus, et al., Repairing a deleterious domestication variant in a floral regulator gene of tomato by base editing. Nat. Genet. 57, 231–241 (2025).

57. A. Abe, et al., Genome sequencing reveals agronomically important loci in rice using MutMap. Nat. Biotechnol. 30, 174–178 (2012).

58. G. Ramstein, C. Moslemi, In vivo validation of predicted fitness effects at single-base resolution in a Brachypodium distachyon mutant population. Zenodo. 10.5281/ZENODO.19145364. Deposited 2026.

