## Supplementary Information for "*In vivo* validation of predicted fitness effects at single-base resolution in a *Brachypodium distachyon* mutant population"

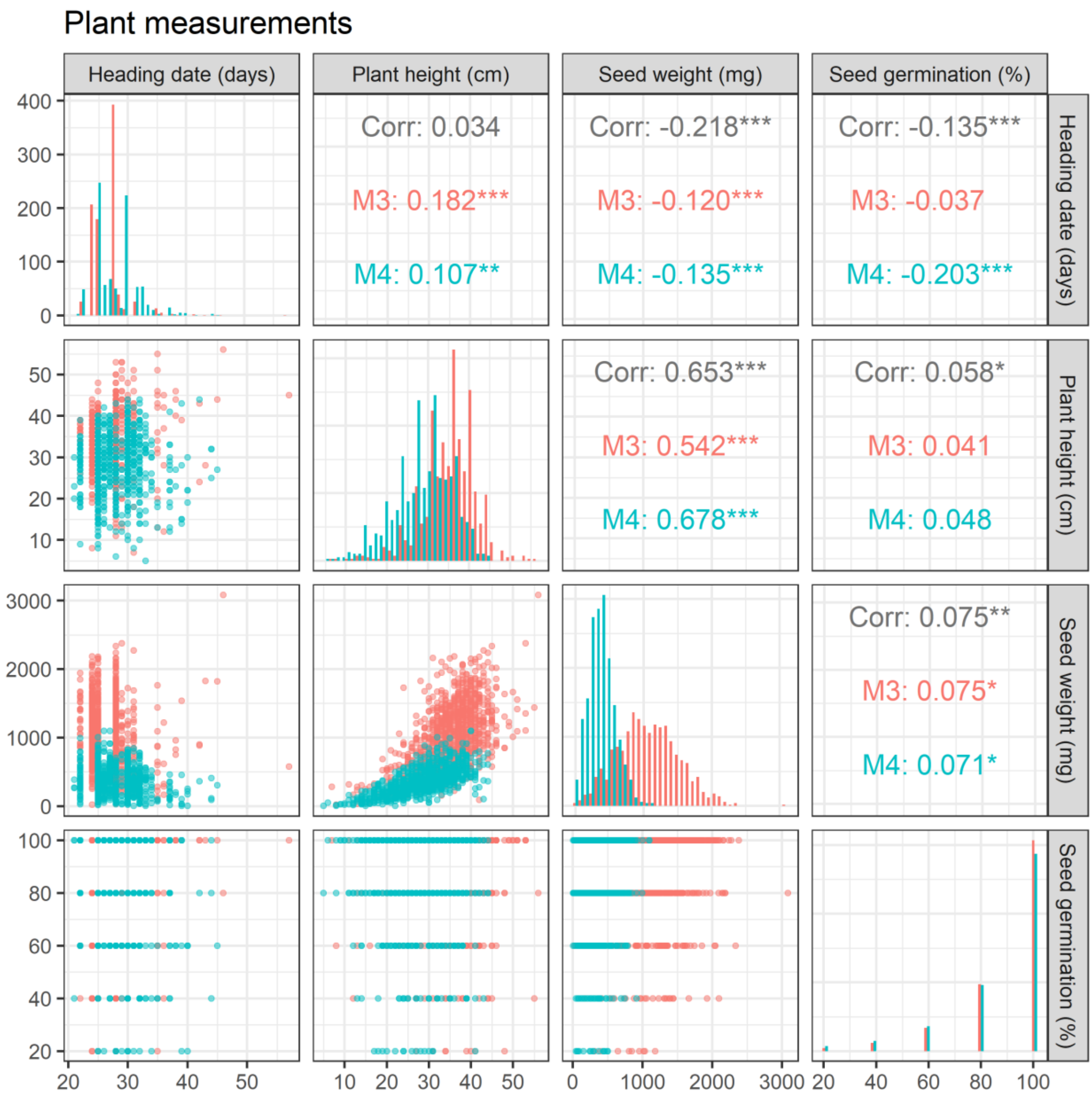


**Figure S1 – Distribution of plant measurements at generation M$_3$ (red) and M$_4$ (blue).** Diagonal: histogram of plant measurements by generation. Below diagonal: pairwise relationship between traits. Above diagonal: Pearson's correlation across and within generations between traits.

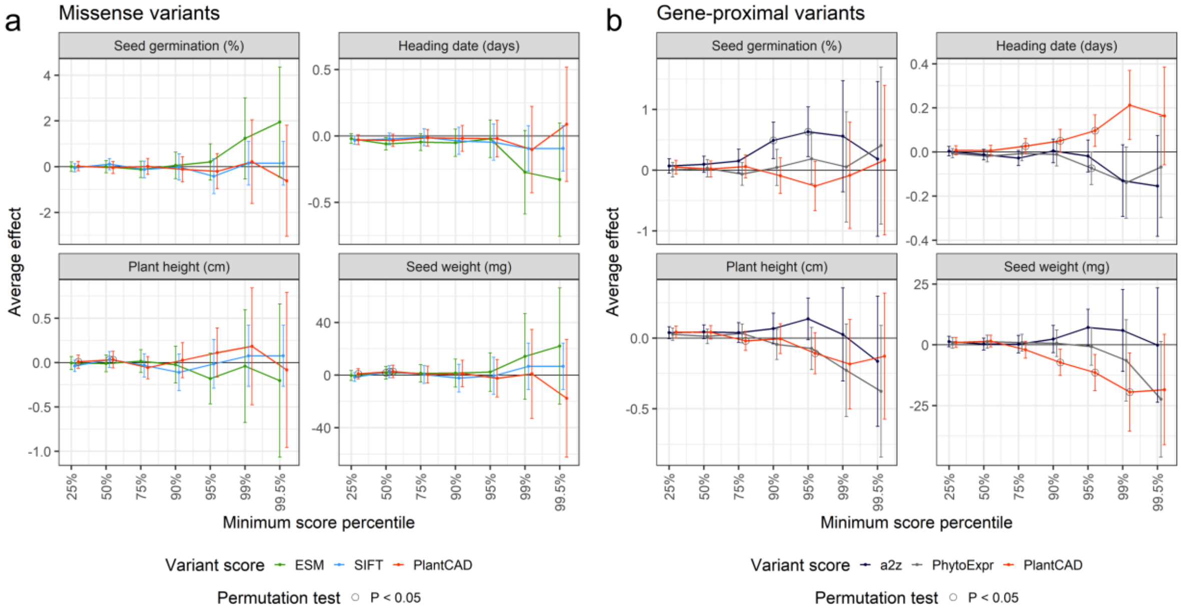


**Figure S2 – Functional enrichment of homozygous variants prioritized for predicted positive effects.** Average effect (point estimate + 95% confidence interval) of homozygous G:C-to-A:T singletons on seed germination, heading date, plant height, and total seed weight at the $M_3$ and $M_4$ generations. Black circles refer to average effects that are significant based on 100 permutations of predicted variant effects (empirical P-value lower than 0.05). (a) Prioritizations of missense variants (variants causing amino-acid change in protein sequences of primary transcripts) by SIFT (multiple sequence alignments), ESM (protein language model), or PlantCAD (genomic language model). (b) Prioritization of gene-proximal variants (variants within 4000 bp of a gene, excluding those in protein-coding regions) by a2z (chromatin-state model), PhytoExpr (sequence-to-expression model), or PlantCAD.
